# Hybphaser identifies hybrid evolution in Australian Thelypterid ferns

**DOI:** 10.1101/2021.11.29.470477

**Authors:** Zoë Bloesch, Lars Nauheimer, Thaís Elias Almeida, Darren Crayn, Ashley Raymond Field

## Abstract

Hybridisation can lead to reproductive isolation and consequently speciation. It has been proposed to play an important role in fern evolution, but has been difficult to investigate. This study explores the utility of target sequence capture and reference guided read phasing to investigate the role of evolutionary reticulation in ferns using Australian Thelypteridaceae as a model. The bioinformatics workflow HybPhaser was used to assess divergence between alleles, phase sequence reads to references to construct accessions resembling parental haplotpes, and include them in phylogenetic and network analyses to detect hybrids and parentage. This approach identified two novel hybrid lineages in Thelypteridaceae, one occurring between two different genera (*Abacopteris* and *Christella*), and provided evidence that reticulation is likely to have played an important role in the diversification of Australian thelypterids. In addition, hybrid phasing successfully reduced conflicting data and improved overall resolution in the Thelypteridaceae phylogeny, highlighting the power of this approach for reconstructing evolutionary history in reticulated lineages.

## 1. Introduction

Speciation resulting from vicariance or dispersal, selection, gene polymorphism and genome duplication are well understood (Pinheiro et al., 2018; Galtier, 2019). There is a growing body of research revealing that hybridization is also an important contributor to speciation, and that it may be more common than previously known (Mallet, 2007; Soltis & Soltis, 2009; Abbott et al., 2013; Vaux et al., 2016; Kinosian et al., 2020; Nauheimer et al., 2020; Requena et al., 2020, Rutherford 2020; Ottenburghs, 2020). A reticulate evolutionary history may be important for generation of future diversification and a key to the successful radiation of some groups (Goulet et al., 2017; Liu et al., 2020; Requena et al., 2020). Hybrid speciation and introgression may be an important reason evolutionary relationships have been difficult to resolve using phylogeny in some groups of plants (Mallet et al., 2016). Identification of the contribution of hybrid evolution is critical to a better understanding of evolutionary relationships (Abbott et al., 2013) and ultimately the conservation of species.

Detecting and investigating hybridisation and reticulation has long been a challenge for phylogenetic studies. The recent use of targeted sequence capture, which retrieves hundreds of conserved loci across a genome and sequences from all haplotypes (Cronn et al., 2012; Breinholt et al., 2020) has the potential to be used to investigate heterozygosity and identify hybrids by phasing. A method called read-backed phasing (separating DNA sequence reads that belong to different copies of loci) was used to investigate heterozygosity on non-hybrid taxa by Kates et al. (2018) who found that heterozygous samples were not a source of conflicting data and concluded that phasing had limited impact on phylogenetic reconstructions. In contrast, Andermann et al. (2019) found that phasing improved the accuracy of phylogenetic tree topology and divergence estimates. Read-backed phasing, however, does not link phased reads according to which haplotype they belong to, so the method cannot determine the presence of, or parentage of a hybrid (Nauheimer et al., 2020). A new approach called reference guided allele phasing phases sequence reads to phylogenetically informed references to construct accessions resembling parental haplotpes providing the ability to infer the presence of and parentage of hybrids (Nauheimer et al. 2020).

Reticulate evolution is considered particularly important in ferns, which in general hybridise more frequently than angiosperms (Haufler et al., 2000, Rothfels et al., 2015). The large global fern family Thelypteridaceae has a history of difficulty to resolve phylogeny and species diversity (see Almeida et al., 2016; Fawcett and Smith, 2021) but the contribution of hybrid speciation and reticulation to diversification of this family has not been investigated. Currently, 1034 species and a range from one to 37 genera have been recognised (Smith, 1990; Almeida et al., 2016; Fawcett and Smith, 2021). Thirteen genera and 24 species occur in Australia (Field, 2020; Fawcett and Smith, 2021) with several populations known that do not correspond to presently circumscribed entities.

Six phylogenetic studies have been undertaken on the Thelypteridaceae to date (Smith and Cranfill, 2002; He and Zhang, 2012; Almeida et al., 2016; Patel et al., 2019; Kuo et al., 2020). These studies have had low phylogenetic sampling and have each used five or fewer Sanger-sequenced loci from the chloroplast genome. Sanger sequencing of chloroplast loci does not readily allow detection and characterisation of heterozygosity, polyploids, hybrids and reticulate evolution. Unidentified conflicting signal in Sanger datasets substantially limits their capacity to infer a well-supported tree (Andermann et al., 2019). To date, no phylogenetic study has focused on Australian Thelypteridaceae species, although some Australian species have been included in previous studies and no study has addressed potential for hybridisation in the family.

Two distinctive taxonomically unassigned Thelypteridaceae have been collected recently in the Australian Wet Tropics. One entity, first recorded at the Russell River is hereafter referred to as Russell River Fern, and the other, first recorded from the Tully River, is hereafter referred to as Tully River Fern (Fig. 1). Russell River Fern is known from three populations spread across the Wet Tropics: in the Daintree region, the Russell River region, and the Hinchinbrook region. It is morphologically similar to *Abacopteris aspera* (C.Presl) Ching and has been previously misidentified as that species but differs in many characters, especially in having lobed pinnae and frond shape similarities to the genus *Christella* H.Lév. (Fig. 1). Tully River Fern is known from a single population on the Tully River. It is morphologically similar to both *Amblovenatum opulentum* (Kaulf.) J.P.Roux and the genus *Christella* (Fig. 1). It was identified as *Amblovenatum tildeniae* (Holttum) T.E.Almeida & A.R.Field (in Field, 2020) but has also been hypothesised to be a hybrid between *Christella dentata* (Forssk.) Brownsey & Jermy and *Christella subpubescens* (Blume) Holttum when originally collected.

**Figure 1.**
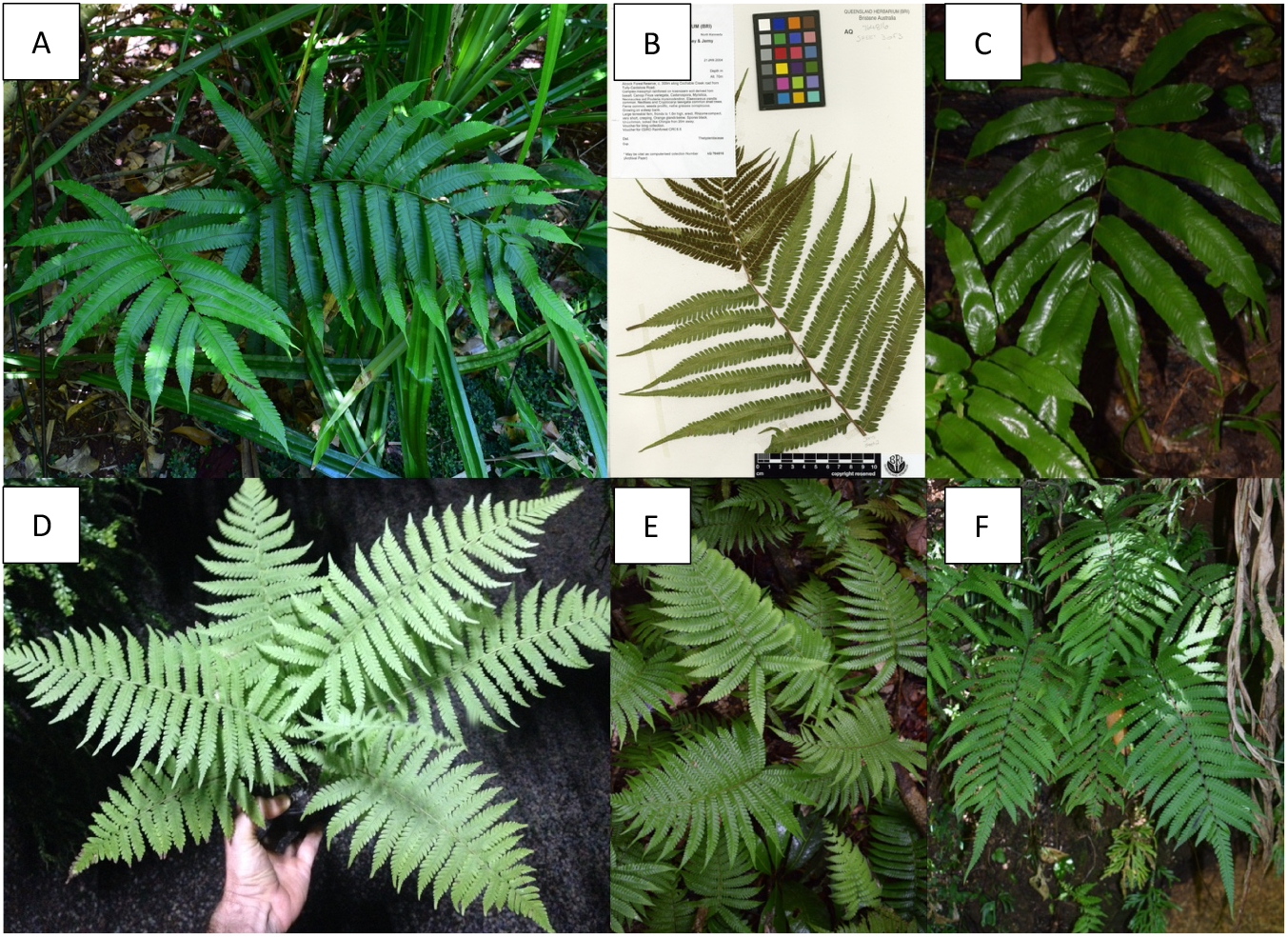
Images of some Thelypteridaceae species and putative novelties. A. Russell River Fern. B. Tully River Fern. C. *Abacopteris aspera*. D. *Amblovenatum queenslandicum*. E. *Christella parasitica*. F. *Christella subpubescens*.

We undertook a hybrid-detecting phylogenomic study of Russell River Fern and Tully River Fern using a recently developed reference guided read phasing method – HybPhaser (Nauheimer et al., 2020) – to assess their putative hybrid origins and determine their parentage and phylogenetic relationships.

## 2. Materials and methods

### 2.1 Dataset preparation

#### 2.1.1 Sampling

The sampling comprised 29 samples of Australian Thelypteridaceae including 20 of 24 species, all 11 genera, both subfamilies recorded for Australia and both putative novelties (Field, 2020). (Suppl. material, Table S1). *Macrothelypteris torresiana* (Gaudich.) Ching is included as an outgroup, based on its phylogenetic placement in the subfamily Phegopteridoideae Salino, A.R.Sm. & T.E.Almeida (Almeida et al., 2016). The remaining samples are from the subfamily Thelypteridoideae, which contains the bulk of the species in the family (Almeida et al., 2016), and two of the four main thelypteroid clades were represented in the cleaned dataset (Almeida et al., 2016). Vouchers are accessioned in the Queensland Herbarium (BRI) and are cited as the primary collector surname and collector number, e.g. *Almeida 4485*.

#### 2.1.2 DNA extraction, library enrichment and sequencing

Total genomic DNA was extracted from all samples using a modified cetyl-trimethylammonium bromide (CTAB) method (Doyle and Doyle, 1987). Sample quality was determined using the Nanodrop Microvolume Spectrophotometer and Qubit Fluorometric Quantification (ThermoFisher Scientific, Waltham, Massachusetts). Library enrichment and sequencing with targeted sequence capture were performed at RAPiD Genomics (Florida, USA), utilising a bait set developed by the GoFlag project which targets 451 nuclear loci of at least 120 base pair length across all flagellate plants (Breinholt et al., 2020).

#### 2.1.3 Trimming and sequence assembly

Sequence reads were trimmed by removing Illumina adapters and low-quality bases and reads using Trimmomatic (Bolger et al., 2014) (illuminaclip 2:30:10, leading 20, trailing 20, sliding window 4:20). Trimmed reads of less than 30 bases were excluded, and samples that recovered less than 10% of the average number of loci recovered were excluded. HybPiper (Johnson et al., 2016) was used to assemble gene sequences including intron regions using the Leptosporangiate-Polypodiales targeted sequences provided by GoFlag. HybPiper performs de novo assembly of reads matching to a single locus to generate contigs that are then connected across exons to generate supercontigs representing gene sequences. The inclusion of intron regions can be used to generate intronerated supercontigs.

### 2.2 Data analysis

#### 2.2.1 Dataset optimisation and assessment of heterozygosity

The bioinformatics workflow, HybPhaser (Nauheimer et al., 2020), was used to detect putative hybrids by assessing the conflicting signal across the dataset and phase sequence reads to generate phased accessions where possible. In a first step, HybPhaser provided the option to optimise the dataset by removing samples or loci with poor sequence recovery. Here loci with less than 20% sequence recovery, and samples with less than 20% of loci recovered or less than 50% of sequence length recovered were removed. Most importantly, HybPhaser provided assessment of divergence between alleles or haplotypes by remapping sequence reads for each sample to the de novo assembled intronerated supercontigs of each locus in order to generate consensus sequences that contain ambiguity codes where single nucleotide polymorphism (SNPs) were encountered. Assessment of SNPs across all loci provided information on allele divergence (AD, % of SNPs across all loci) and locus heterozygosity (LH, % of loci with SNPs). Hybrids contain alleles from divergent parental species and thus are expected to have high proportions of LH as almost all loci have some divergence and have an AD that resembles the divergence between the parents (Nauheimer et al., 2020). In addition, the assessment of SNPs across loci allows to detect and remove putative paralogs, i.e. loci that have unusual high amount of SNPs compared to other loci. Here loci for all samples were removed that had outlying values for the mean proportion of SNPs across all samples and subsequently all loci were removed for each sample individually that had an outlier value for the proportion of SNPs compared to other loci. Finally, sequence lists were generated for each locus containing all samples, as well as for all samples containing all loci using consensus sequences. The software PRANK (Löytynoja, 2014) was used to align the sequence list of each locus. Columns with more than 50% of gaps were removed using TrimAl (Capella-Gutierrez et al., 2009) to reduce the proportion of missing data.

#### 2.2.2 Phylogenetic analysis

To ensure a robust phylogenetic reconstruction of the relationship between samples, three analyses were performed: a supermatrix analysis of the concatenated genes, multi-species coalescence analysis (MSC) gene tree summary phylogeny, and a network reconstruction. Maximum likelihood (ML) analyses of supermatrices are widely used, being relatively fast, and an improvement on simpler and computationally faster parsimony-based methods, because they address evolutionary rate variation among lineages by incorporating a substitution model (Nguyen et al., 2015). However, if incomplete lineage sorting or divergent gene evolution is present, gene trees will show different evolutionary histories from the species tree, and this cannot be detected using ML analysis of a concatenated dataset (Roch et al., 2019). Multi-species coalescence (MSC) generates a summary tree from ML gene trees and can show discordance between gene trees due to incomplete lineage sorting (Mirarab et al., 2014). However, neither ML nor MSC methods can effectively deal with reticulation, so network reconstruction was also used, which can visualise reticulation better than the other two methods (Huson and Bryant, 2006). Since no single method gives a complete picture, all three were used together in this study. IQ-TREE2 (Minh et al., 2020b) was used to perform a ML analysis of the concatenated gene dataset with the edge-partitioned substitution model approach and ultrafast bootstrap (1000 replicates) (Hoang et al., 2018). IQ-TREE2 was used to generate gene concordance factors (gCF), which give the percentage of informative gene trees containing the same branch, and site concordance factors (sCF), which give the percentage of informative alignment sites that support the branch (Minh et al., 2020a). ASTRAL III (Zhang et al., 2018) was used to generate a summary phylogeny tree based on ML gene trees generated with IQ-TREE2 and ModelFinder (Kalyaanamoorthy, 2017) to determine the best substitution model for each gene. SplitsTree5 (Huson and Bryant, 2006) was used to infer a split network from the concatenated alignment using Neighbour Net and the Splits Network Algorithm with Equal Angle and Convex Hull.

##### Clade association

Accessions of hybrids with parents from divergent clades contain reads that associate with references from these divergent clades and thus can be detected and sorted accordingly. Ten references were chosen that represented major clades in the phylogenies (supermatrix and gene tree summary phylogeny) and that had high sequence coverage but low LH and AD. HybPhaser used BBSplit (BBMap, v38.47) to simultaneously map reads to all clade references simultaneously and to record the proportions of reads that mapped unambiguously to only one reference.

##### Phasing

Samples with reads mapping in high proportions to multiple clade references and with high LH and AD were selected for phasing as they represent putative hybrid accessions with parents from the clades represented by those references. Again, HybPhaser was used to map reads with BBSplit to multiple references simultaneously, but only to the references with high read association. Reads were saved in separate files according to their association. Reads matching unambiguously to one reference were saved in a separate read file, while reads matching similarly well to multiple references were saved in all read files. Each phased read file theoretically represented one of the haplotypes that a hybrid plant inherited from its parents.

The read files of the phased accessions were processed similar to the original read files. First, HybPiper was used for assembly, generating supercontigs including intron regions. Then HybPhaser was used for dataset optimisation with the same settings as above for the phased accessions. The same loci that were removed as putative paralogs across all samples in the original dataset were removed from the phased accessions. Similarly, all outlier loci for each individual phased accession were removed. Sequence lists for the phased accessions were generated in HybPhaser and combined with the original dataset while removing the non-phased accessions of the samples that were phased.

Finally, the resulting sequence lists including phased accessions were aligned and trimmed to generate alignments and phylogenetic analyses were performed exactly as for the non-phased dataset to generate a supermatrix tree, a gene tree summary phylogeny and a network.

## 3. Results

### 3.1 Dataset assembly

On average 365.75 out of the 451 target loci were recovered, and on average 83.56% of the targeted sequence length was recovered for each sample (Suppl. material, Table S3). Two separate assemblies: exons only and exons + introns were prepared. The assemblies of exons + introns (i.e. target loci (exons) and additional non-target loci (introns)) were chosen for analysis because they included 3.3x more data than those with exons only.

The cleaning step removed one failed locus, leaving 407 loci with high recovery across samples. It removed 49 loci with >1.5% SNPs across all samples (putative paralogs or the result of contamination or sequencing error) (Suppl. material, Fig. S1). It also removed an average of 12.48 loci from each sample, using the same threshold (Suppl. Material, Fig. S1, Fig. S2 and Table S4). After cleaning, the combined exons + introns sequences were 237,644 base pairs long.

### 3.2 Assessment of heterozygosity and reference guided allele phasing

Around half of the accessions had moderate locus heterozygosity (>50% of loci with >0% SNPs) and allele divergence (>0.5%), and ten had high locus heterozygosity (>90% of loci with >0% SNPs) and high allele divergence (≥1%) including Russell River Fern and Tully River Fern (Table 1). High locus heterozygosity was generally associated with high allele divergence, and Russell River Fern was an outlier with very high allele divergence (Fig. 2). *Amblovenatum queenslandicum* (Holttum) T.E.Almeida & A.R.Field, *Reholttumia costata* S.E.Fawc. & A.R.Sm.*, Chingia australis* Holttum and *Amblovenatum terminans* (Panigrahi) J.P.Roux all had low locus heterozygosity (<30% of loci with >0% SNPs) and low allele divergence (<0.13%).

**Table 1.**
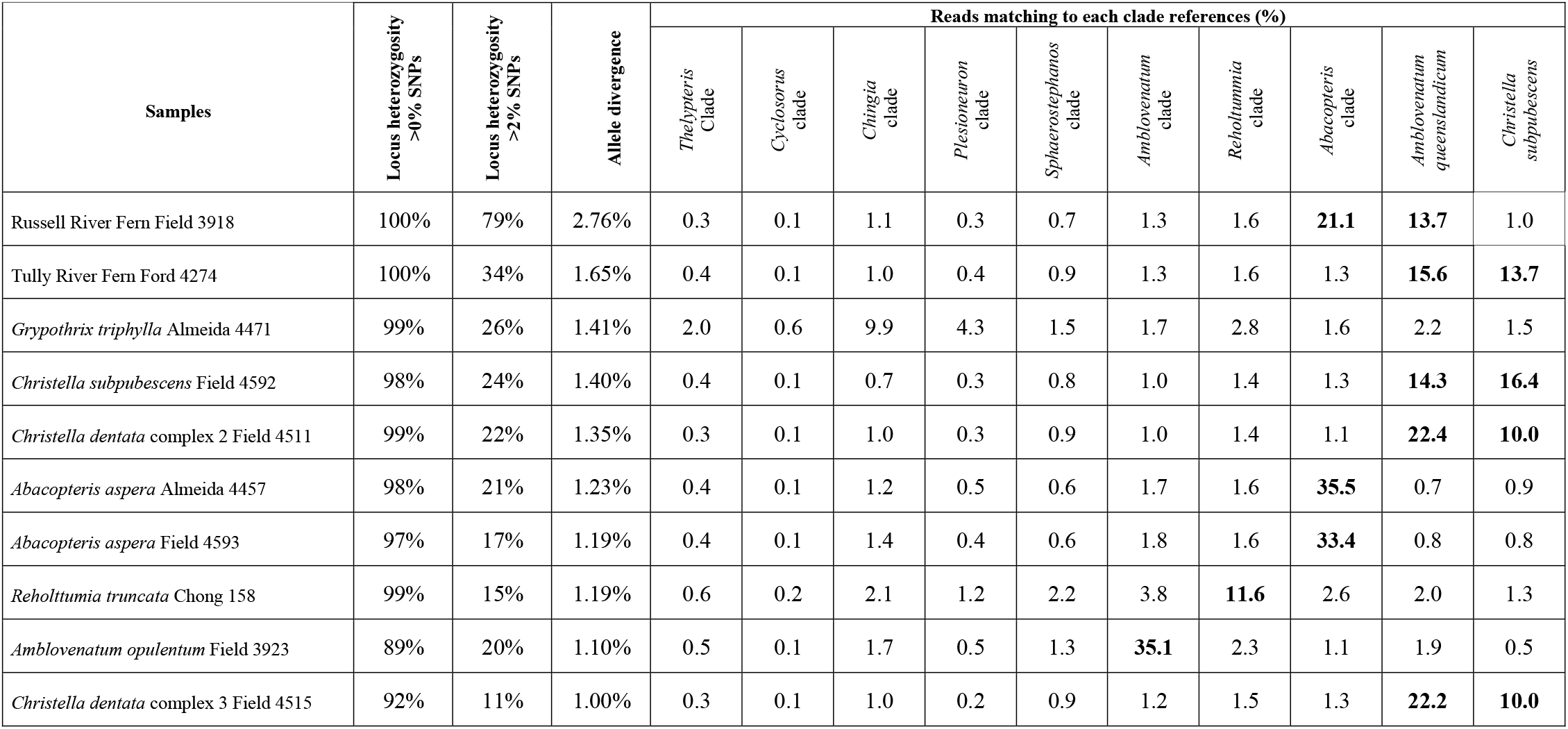
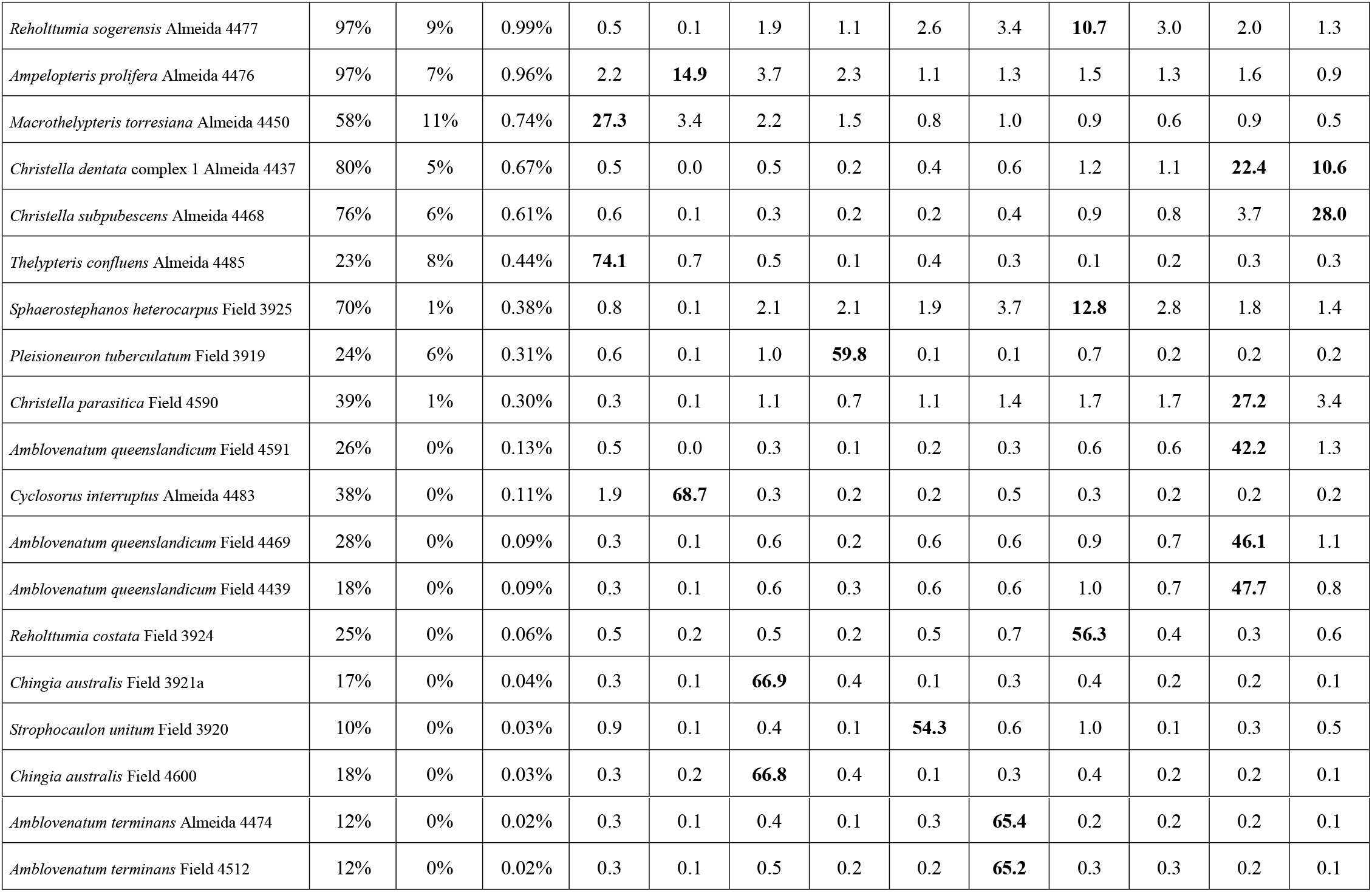
Mean locus heterozygosity (showing the proportion of loci containing >0% and >2% SNPs) and allele divergence (the proportion of SNPs for each sample), and results of clade association analysis, showing the proportion of reads that matched unambiguously to each clade reference.

| Samples | Locus heterozygosity<br>>0% SNPs | Locus heterozygosity<br>>2% SNPs | Allele divergence | Reads matching to each clade references (%) |  |  |  |  |  |  |  |  |  |
| --- | --- | --- | --- | --- | --- | --- | --- | --- | --- | --- | --- | --- | --- |
|  |  |  |  | *Thelypteris*<br>Clade | *Cyclosorus*<br>clade | *Chingia*<br>clade | *Plesioneuron*<br>clade | *Sphaerostephanos*<br>clade | *Amblovenatum*<br>clade | *Reholtumia*<br>clade | *Abacopteris*<br>clade | *Amblovenatum*<br>*queenslandicum* | *Christella*<br>*subpubescens* |
| Russell River Fern Field 3918 | 100% | 79% | 2.76% | 0.3 | 0.1 | 1.1 | 0.3 | 0.7 | 1.3 | 1.6 | **21.1** | **13.7** | 1.0 |
| Tully River Fern Ford 4274 | 100% | 34% | 1.65% | 0.4 | 0.1 | 1.0 | 0.4 | 0.9 | 1.3 | 1.6 | 1.3 | **15.6** | **13.7** |
| *Grypothrix triphylla* Almeida 4471 | 99% | 26% | 1.41% | 2.0 | 0.6 | 9.9 | 4.3 | 1.5 | 1.7 | 2.8 | 1.6 | 2.2 | 1.5 |
| *Christella subpubescens* Field 4592 | 98% | 24% | 1.40% | 0.4 | 0.1 | 0.7 | 0.3 | 0.8 | 1.0 | 1.4 | 1.3 | **14.3** | **16.4** |
| *Christella dentata* complex 2 Field 4511 | 99% | 22% | 1.35% | 0.3 | 0.1 | 1.0 | 0.3 | 0.9 | 1.0 | 1.4 | 1.1 | **22.4** | **10.0** |
| *Abacopteris aspera* Almeida 4457 | 98% | 21% | 1.23% | 0.4 | 0.1 | 1.2 | 0.5 | 0.6 | 1.7 | 1.6 | **35.5** | 0.7 | 0.9 |
| *Abacopteris aspera* Field 4593 | 97% | 17% | 1.19% | 0.4 | 0.1 | 1.4 | 0.4 | 0.6 | 1.8 | 1.6 | **33.4** | 0.8 | 0.8 |
| *Reholtumia truncata* Chong 158 | 99% | 15% | 1.19% | 0.6 | 0.2 | 2.1 | 1.2 | 2.2 | 3.8 | **11.6** | 2.6 | 2.0 | 1.3 |
| *Amblovenatum opulentum* Field 3923 | 89% | 20% | 1.10% | 0.5 | 0.1 | 1.7 | 0.5 | 1.3 | **35.1** | 2.3 | 1.1 | 1.9 | 0.5 |
| *Christella dentata* complex 3 Field 4515 | 92% | 11% | 1.00% | 0.3 | 0.1 | 1.0 | 0.2 | 0.9 | 1.2 | 1.5 | 1.3 | **22.2** | **10.0** |
| *Reholtumia sogerensis* Almeida 4477 | 97% | 9% | 0.99% | 0.5 | 0.1 | 1.9 | 1.1 | 2.6 | 3.4 | **10.7** | 3.0 | 2.0 | 1.3 |
| *Amplopteris prolifera* Almeida 4476 | 97% | 7% | 0.96% | 2.2 | **14.9** | 3.7 | 2.3 | 1.1 | 1.3 | 1.5 | 1.3 | 1.6 | 0.9 |
| *Macrothelypteris torresiana* Almeida 4450 | 58% | 11% | 0.74% | **27.3** | 3.4 | 2.2 | 1.5 | 0.8 | 1.0 | 0.9 | 0.6 | 0.9 | 0.5 |
| *Christella dentata* complex 1 Almeida 4437 | 80% | 5% | 0.67% | 0.5 | 0.0 | 0.5 | 0.2 | 0.4 | 0.6 | 1.2 | 1.1 | **22.4** | **10.6** |
| *Christella subpubescens* Almeida 4468 | 76% | 6% | 0.61% | 0.6 | 0.1 | 0.3 | 0.2 | 0.2 | 0.4 | 0.9 | 0.8 | 3.7 | **28.0** |
| *Thelypteris confluens* Almeida 4485 | 23% | 8% | 0.44% | **74.1** | 0.7 | 0.5 | 0.1 | 0.4 | 0.3 | 0.1 | 0.2 | 0.3 | 0.3 |
| *Sphaerostephanos heterocarpus* Field 3925 | 70% | 1% | 0.38% | 0.8 | 0.1 | 2.1 | 2.1 | 1.9 | 3.7 | **12.8** | 2.8 | 1.8 | 1.4 |
| *Pleisoneuron tuberculatum* Field 3919 | 24% | 6% | 0.31% | 0.6 | 0.1 | 1.0 | **59.8** | 0.1 | 0.1 | 0.7 | 0.2 | 0.2 | 0.2 |
| *Christella parasitica* Field 4590 | 39% | 1% | 0.30% | 0.3 | 0.1 | 1.1 | 0.7 | 1.1 | 1.4 | 1.7 | 1.7 | **27.2** | 3.4 |
| *Amblovenatum queenslandicum* Field 4591 | 26% | 0% | 0.13% | 0.5 | 0.0 | 0.3 | 0.1 | 0.2 | 0.3 | 0.6 | 0.6 | **42.2** | 1.3 |
| *Cyclosorus interruptus* Almeida 4483 | 38% | 0% | 0.11% | 1.9 | **68.7** | 0.3 | 0.2 | 0.2 | 0.5 | 0.3 | 0.2 | 0.2 | 0.2 |
| *Amblovenatum queenslandicum* Field 4469 | 28% | 0% | 0.09% | 0.3 | 0.1 | 0.6 | 0.2 | 0.6 | 0.6 | 0.9 | 0.7 | **46.1** | 1.1 |
| *Amblovenatum queenslandicum* Field 4439 | 18% | 0% | 0.09% | 0.3 | 0.1 | 0.6 | 0.3 | 0.6 | 0.6 | 1.0 | 0.7 | **47.7** | 0.8 |
| *Reholtumia costata* Field 3924 | 25% | 0% | 0.06% | 0.5 | 0.2 | 0.5 | 0.2 | 0.5 | 0.7 | **56.3** | 0.4 | 0.3 | 0.6 |
| *Chingia australis* Field 3921a | 17% | 0% | 0.04% | 0.3 | 0.1 | **66.9** | 0.4 | 0.1 | 0.3 | 0.4 | 0.2 | 0.2 | 0.1 |
| *Strophocaulon unitum* Field 3920 | 10% | 0% | 0.03% | 0.9 | 0.1 | 0.4 | 0.1 | **54.3** | 0.6 | 1.0 | 0.1 | 0.3 | 0.5 |
| *Chingia australis* Field 4600 | 18% | 0% | 0.03% | 0.3 | 0.2 | **66.8** | 0.4 | 0.1 | 0.3 | 0.4 | 0.2 | 0.2 | 0.1 |
| *Amblovenatum terminans* Almeida 4474 | 12% | 0% | 0.02% | 0.3 | 0.1 | 0.4 | 0.1 | 0.3 | **65.4** | 0.2 | 0.2 | 0.2 | 0.1 |
| *Amblovenatum terminans* Field 4512 | 12% | 0% | 0.02% | 0.3 | 0.1 | 0.5 | 0.2 | 0.2 | **65.2** | 0.3 | 0.3 | 0.2 | 0.1 |

**Figure 2.**
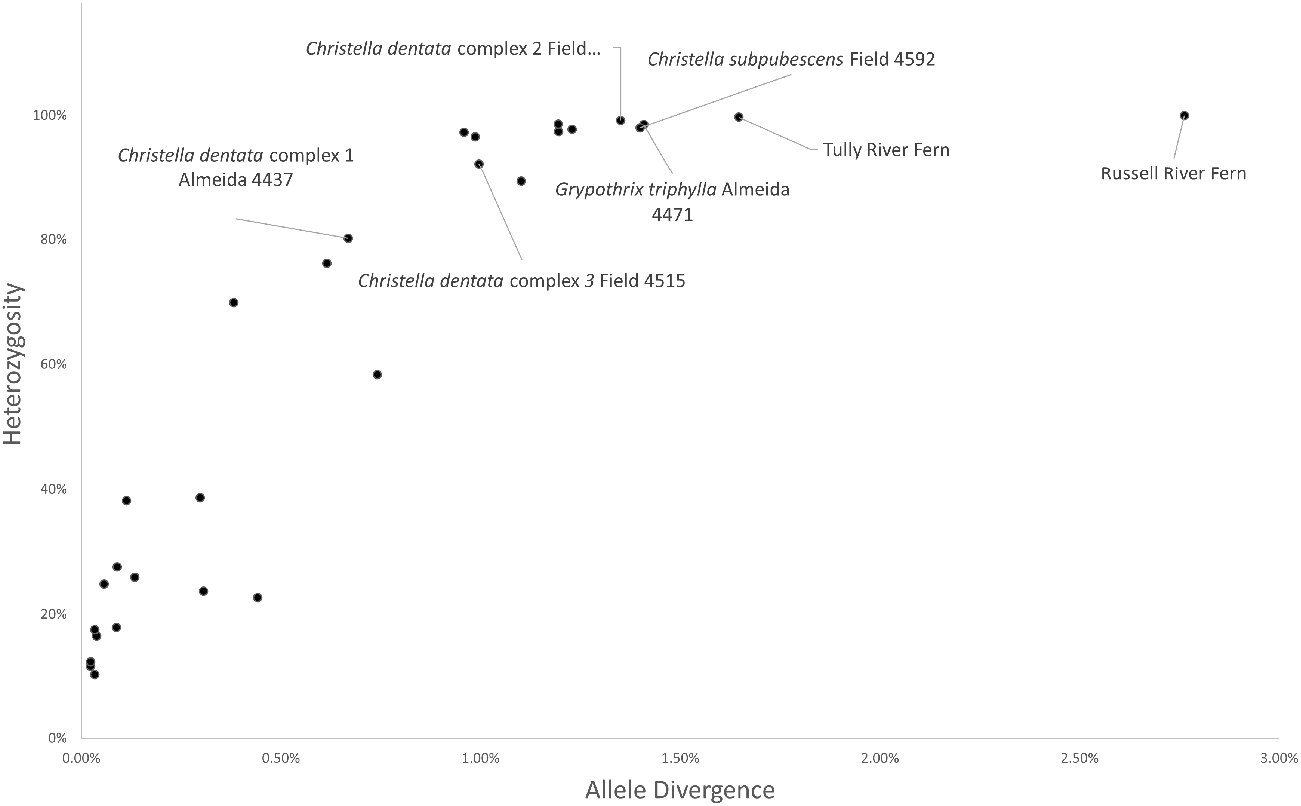
Scatterplot showing the heterozygosity and allele divergence for all samples. Putative hybrid samples are labelled.

Of the samples with high heterozygosity and allele divergence, six had reads which mapped unambiguously and in high proportions to two reference clades, therefore indicating that these were putative hybrids. These were the Russell River Fern, Tully River Fern, *Christella subpubescens* Field 4592, *Christella dentata* complex 2 Field 4511, *Christella dentata* complex 3 Field 4515 and *Christella dentata* complex 1 Almeida 4437. Reads of the three *Christella dentata* complex accessions phased to two clades in a strict 2:1 ratio, while Russell River Fern phased in a 2:3 ratio, and the other hybrids phased in a 1:1 ratio (Table 1). For the phased accession, all samples were recovered, and 40 failed loci were removed. On average 10.91 loci with a high proportion of SNPs were removed from each sample. After cleaning there were 35 samples (putative hybrids were replaced with two phased samples each). As with the unphased dataset, the assemblies of exons + introns were chosen for analysis because they included 2.47x more data than exons only (197,446 base pairs for exons + introns).

### 3.3 Phylogenetic and network analysis

#### 3.3.1 Data before phasing

ML analysis of the data prior to reference guided allele phasing resolved most nodes with moderate to strong support (Fig. 3). *Thelypteris confluens* (Thunb.) C.V.Morton (*Thelypteris* clade) was placed sister to the Cyclosoroid clade which includes all the other accessions in this study. Within the Cyclosoroid clade, *Ampelopteris prolifera* (Retz.) Copel. and *Cyclosorus interruptus* (Willd.) H.Ito were placed with high support as sister to the Christelloid clade, similarly to Almeida et al. (2016). *Reholttumia* S.E.Fawc. & A.R.Sm. and *Christella* were paraphyletic. *Grypothrix triphylla* (Sw.) S.E.Fawc. & A.R.Sm. is sister to *Plesioneuron* (Holttum) Holttum + *Chingia* Holttum and *Abacopteris aspera* sister to *Christella. Amblovenatum* J.P.Roux comprises two divergent lineages. *Amblovenatum opulentum* (Kaulf.) J.P.Roux and *A. terminans* were sister to *Reholttumia* + *Sphaerostephanos* + *Strophocaulon* whereas *A. queenslandicum* was embedded in *Christella*.

**Figure 3.**
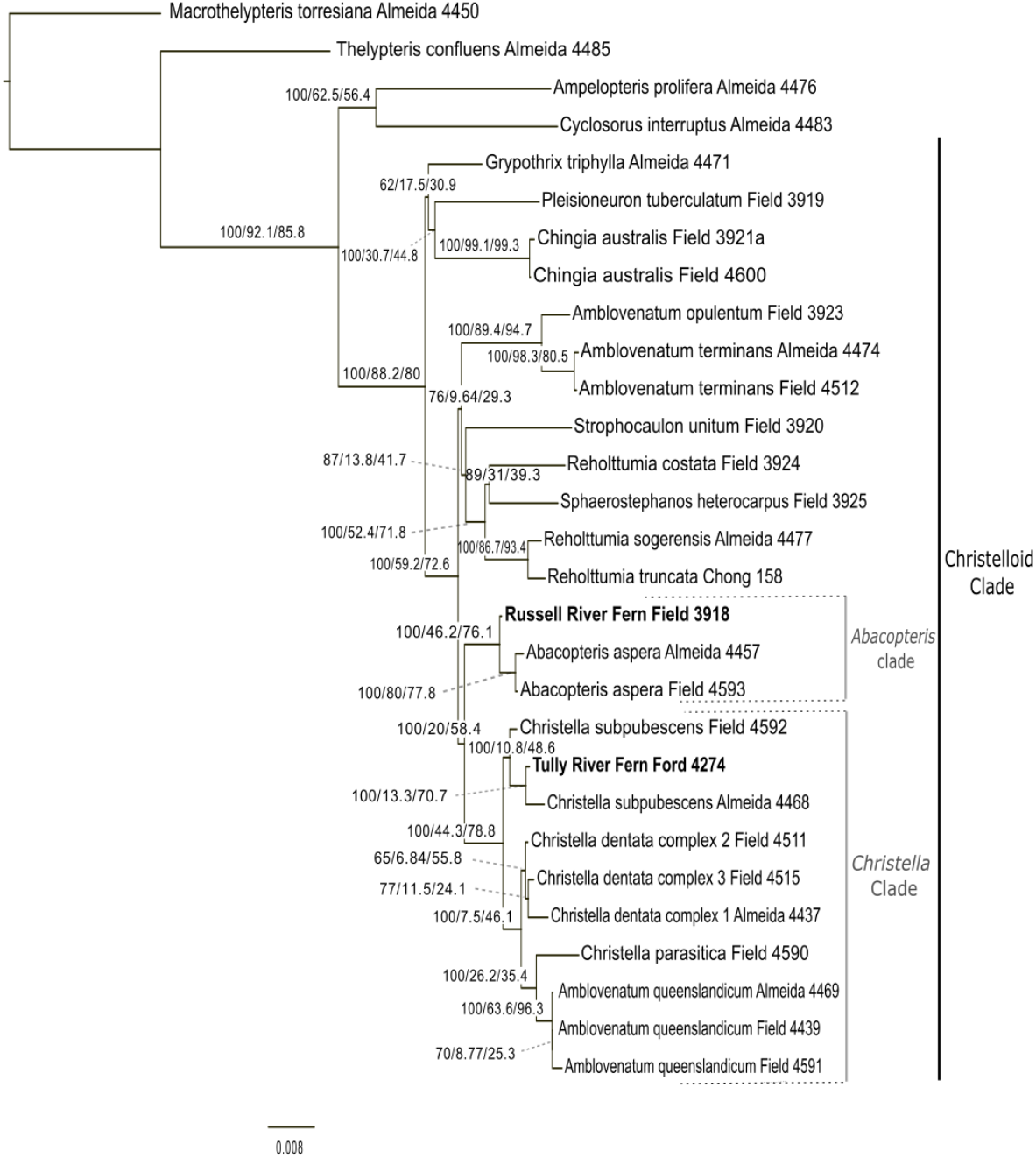
Phylogeny of 29 Thelypteridaceae samples estimated by maximum likelihood analysis of the supermatrix of 363 concatenated nuclear exons and introns. Putative hybrids are coloured in blue and red. Branch support values are ultrafast bootstrap, gene concordance factor and site concordance factors, respectively.

The gene tree summary (Suppl. Material – Figs S3, S4) showed very similar topologies to the supermatrix trees, except for the clade that included *Grypothrix triphylla, Plesioneuron tuberculatum* (Ces.) Holttum and *Chingia australis* as sister to the rest of the Christelloid clade, which had low support (local posterior probability) in the gene tree summary, but high support in the supermatrix tree. This indicates that there was conflict between gene trees (Sayyari and Mirarab, 2016). Other branches were well supported, except within the *Christella* clade.

The split network analysis (Fig. 4) indicated that the data support a genetic relationship between Russell River Fern and both the *Abacopteris* and *Christella* clades. The network also showed substantial reticulation within the *Christella* clade.

**Figure 4.**
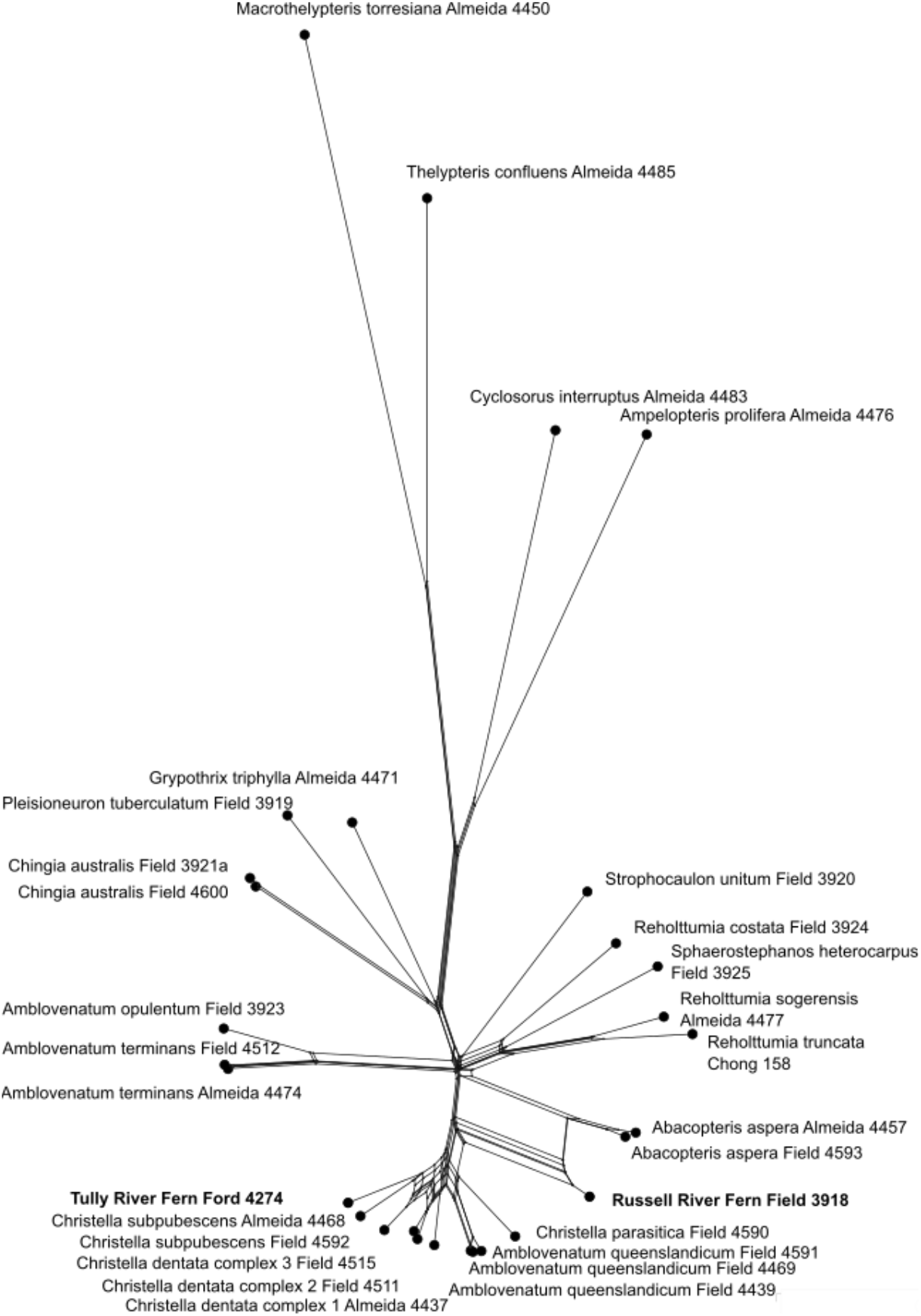
Split network of 29 Thelypteridaceae samples generated by SplitsTree5 analysis (with Neighbour Net settings) of 363 nuclear exons and introns (alleles not phased). End nodes represent samples and edges represent evolutionary relationships. Putative hybrids are bold.

*Grypothrix triphylla* and *Plesioneuron tuberculatum* group with *Chingia australis* as sister to the rest of the Christelloid clade but with low support (Fig. 3), and they were placed differently in the gene summary tree, also with low support.

The Russell River Fern was placed in the *Abacopteris* clade as sister to the two *Abacopteris aspera* accessions, with moderate support. The *Christella* clade was recovered with moderate support but many of its internal nodes were poorly supported (Fig. 3). Support within the *Christella* clade was generally low, and some branches had a bootstrap value of 100 but low gCF and sCF (e.g., 100/11.5/46.1), while some had low support for all measures (e.g., 65/6.84/55.8). Two branches had sCF values below 33% so at least one of the alternative topologies is better supported according to the sCF (Minh et al., 2020). The Tully River Fern was embedded within *Christella subpubescens* as sister to *Christella subpubescens* Almeida 4468 with moderate support (Fig. 3).

#### 3.3.2 Phased data

When phased accessions were included in the supermatrix analysis, they were placed as sister to, or within the same clade as the clade references to which they were mapped (Fig. 5). The two phased accessions of Russell River Fern were placed in different genera with high support, within *Abacopteris aspera* (sister to *A. aspera* Almeida 4457) and sister to the *Amblovenatum queenslandicum* accessions, respectively. The two phased accessions of Tully River Fern were both placed in the *Christella* clade, one as sister to *Christella subpubescens* Almeida 4468, and one as sister to *Christella parasitica* Field 4590 but both with low support. The phased *Christella subpubescens* accessions were both placed in the *Christella* clade, sister to the other *C. subpubescens* sample and *A. queenslandicum*. The *Christella dentata* complex phased accessions converge in two groups within the *Christella* clade. Support values were generally much higher in the phylogenetic tree with phased accessions (Fig. 5) compared to the tree before phasing (Fig. 3), although support within the *Christella* clade remained low.

**Figure 5.**
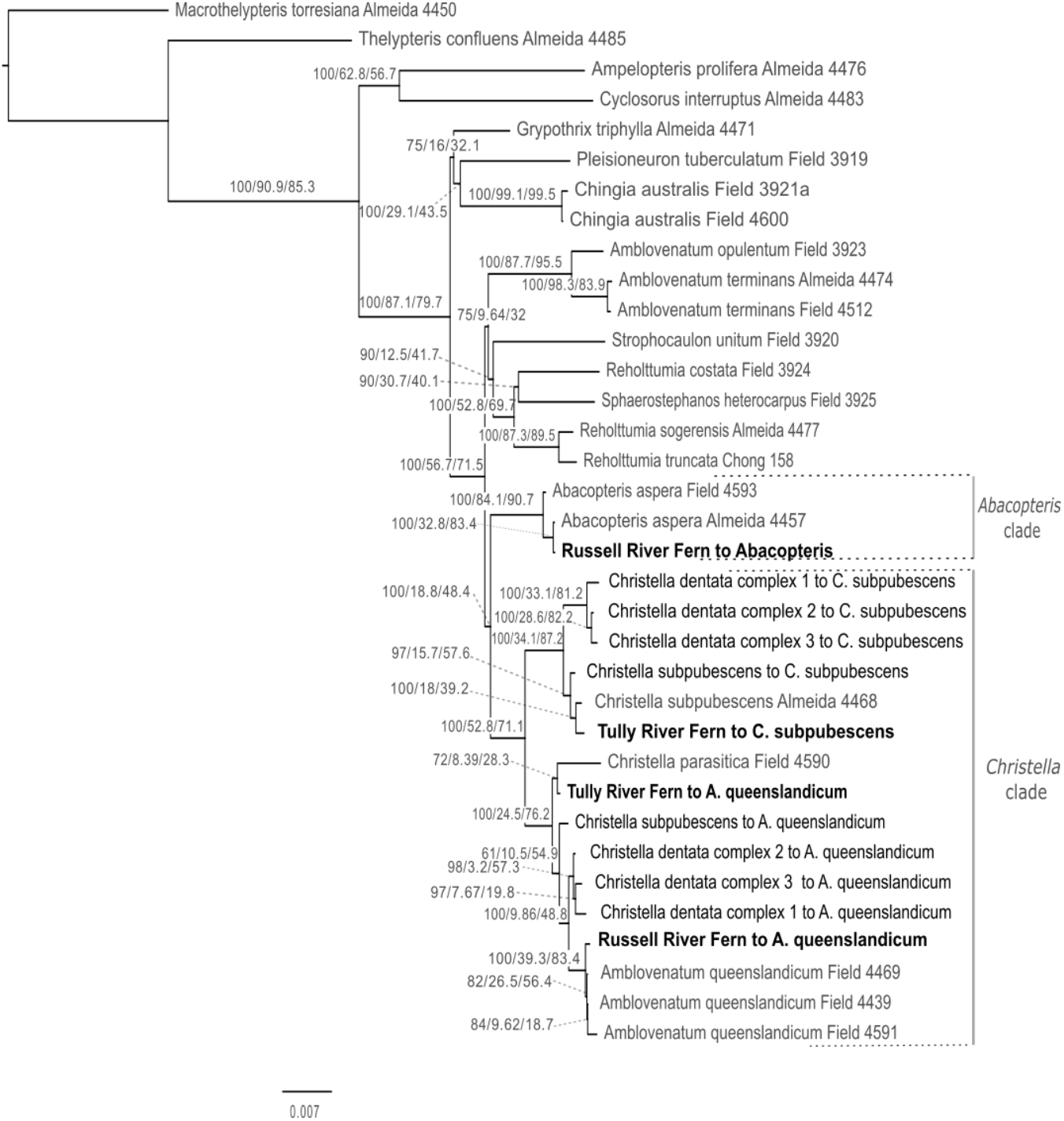
Phylogeny of 29 Thelypteridaceae samples estimated by maximum likelihood analysis of the supermatrix of 363 concatenated 363 nuclear exons and introns. Six putative hybrids are represented by two phased haploid samples each highlighted in bold. Abbreviations (from Suppl. Material, Table S2) indicate which clade reference each sample was phased to. Branch support values are ultrafast bootstrap, gene concordance factor and site concordance factors, respectively.

In the split network with phased samples (Fig. 6), the *Abacopteris* and *Christella* clades were more clearly distinct from each other. Although there was still reticulation within the *Christella* clade, the two subgroups of *Christella* (which can be seen in ML tree, Fig. 3) were also more clearly distinct. These results suggest that phasing alleles reduced conflicting signal in the dataset.

**Figure 6.**
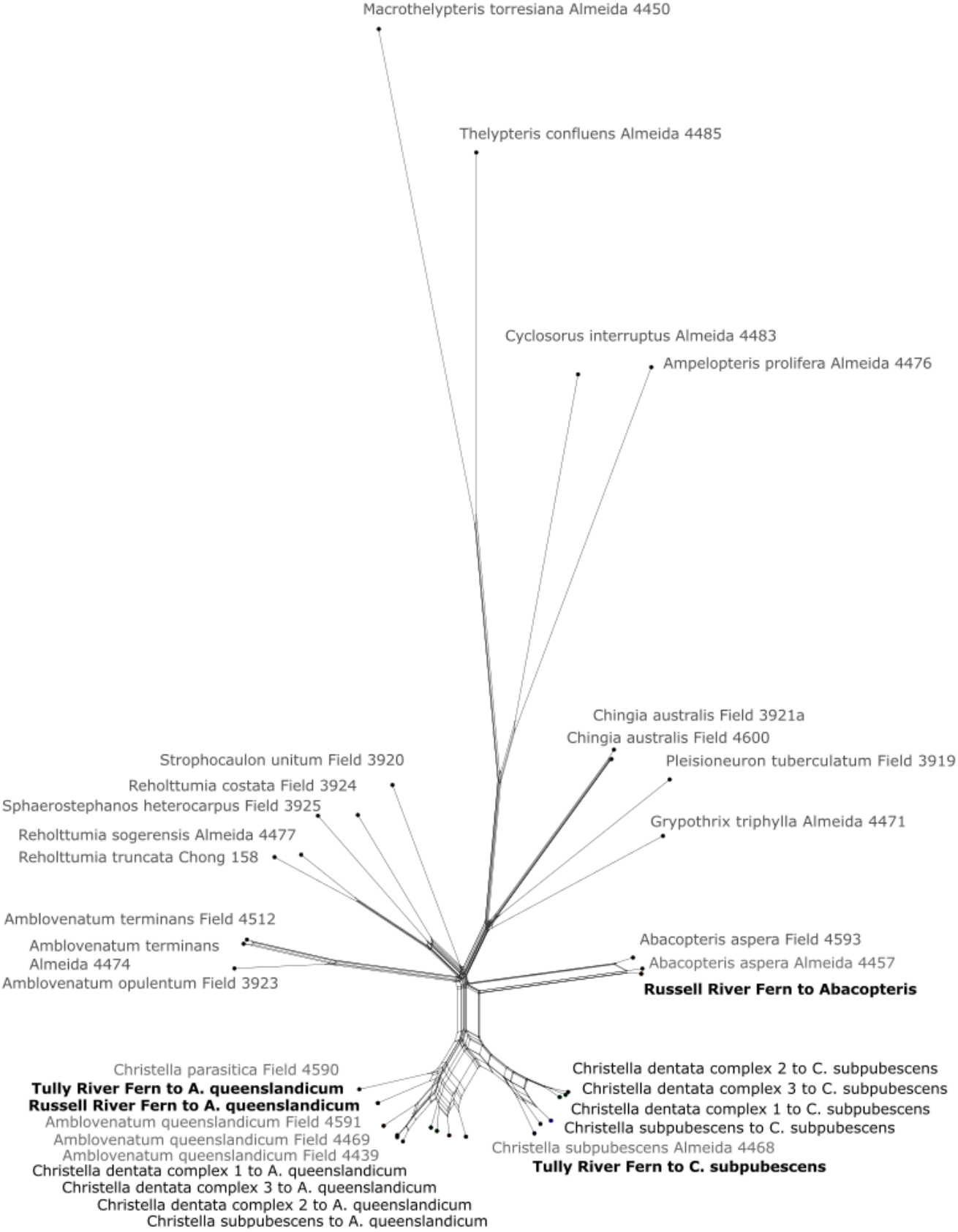
Split network of 29 Thelypteridaceae samples generated by SplitsTree5 analysis (using Neighbour Net settings) of 363 nuclear exons and introns. Six putative hybrids are represented by two phased haploid samples each highlighted in black and bold. End nodes represent samples and edges represent evolutionary relationships.

## 4. Discussion

This is the first molecular phylogenetic study of species of Australian Thelypteridaceae, and the first to use HybPhaser to investigate hybridisation in ferns. The analyses provided strong evidence that both Russell River Fern and Tully River Fern are of hybrid origin and identified the parental lineages. Additionally, the HybPhaser reference guided allele phasing approach identified four previously unrecognised hybrids in the family and showed that there is significant conflicting signal within the *Christella* clade, probably due to reticulate evolution. The results highlight the power of analysing phylogenomic datasets with a hybrid phasing approach to resolve gene and species relationships in reticulated groups.

Russell River Fern showed much higher heterozygosity and allele divergence than any other sample in the dataset, consistent with it being of hybrid origin between diverged parents (Nauheimer et al., 2020). This is confirmed by the placement of the phased haplotype samples in the *Abacopteris* and *Christella* clades with high support (Fig. 5). Allele ratios are theoretically reflected in clade association proportions (Nauheimer et al., 2020) and therefore a 1:1 ratio is expected for diploid first generation (F1) hybrids between diploid parents. Russell River Fern reads phased to the *Abacopteris* and *Christella* clades in approximately a 3:2 ratio, the same as found for one hybrid in the model analysis (Nauheimer et al., 2020). This could indicate a complex ploidy level (Nauheimer et al., 2020), incomplete lineage sorting (Mallet et al., 2016), or possibly a hybridisation event that occurred more than one generation ago. The phased haplotype accessions are placed as sister to *Abacopteris aspera* and *Amblovenatum queenslandicum* with high support. In this case, phasing and phylogenetic analysis gives a clear result and provides strong evidence that these two species are the parents of the Russell River Fern.

Infertility is commonly observed in hybrids (Hornych and Ekrt, 2017) and is a factor in determining whether a hybrid population will persist as a species. Populations of the Russell River Fern have been stable for over 30 years and although they are rarely sexually fertile, they frequently reproduce vegetatively by a long running rhizome (pers. obs.). They are vigorous and have persisted well after disturbances including severe tropical cyclones Winifred (1986), Larry (2006) and Yasi (2011). More work on population size, frequency of hybridisation (i.e., whether all three populations have a single or separate origins), and sexual fertility is required to determine how the conservation of this entity should be managed and its status as a species.

Russell River Fern is the first genetically confirmed inter-generic fern hybrid recorded for Australia. Hybridisation between genera is known to occur occasionally in ferns, and Liu et al. (2020) list twelve confirmed and three possible nothogenera in ferns and lycophytes. A fern nothogenus between genera that diverged 60 million years ago is the most extreme example known (Rothfels et al., 2015).

Tully River Fern had the second highest locus heterozygosity and allele divergence in the analysis (Table 1, Fig. 2) supporting the hypothesis that it is of hybrid origin, from parents that are different but not widely diverged species (Nauheimer et al., 2020). For Tully River Fern, phasing assigned reads in a 1:1 ratio, both to the *Christella* clade reference. This ratio may be evidence it is an F1 hybrid, which is consistent with the single location that may even have resulted from a single recent hybridisation event. Although it is closest to *Christella subpubescens* and *Christella parasitica* (L.) H.Lév., the association of the phased haplotype accessions in the phylogeny were uncertain (support values are very low, Fig. 5). There are two potentials reasons that the parents could not be identified through phasing: firstly, the parents might not have been included in this analysis (*Christella arida* or an unidentified *Christella dentata* complex), and secondly, relationships within the *Christella* clade are generally uncertain and internal support values are very low, even after phasing. In contrast to Russell River Fern, Tully River Fern is thought to reproduce only by spore and does not develop a large localised long-lived clonal colony. More work is required to determine whether the single known occurrence of several plants is the result of one, or multiple, localised hybridisation events, how it should be managed in conservation, and its status as a species. Additionally, more comprehensive sampling of the *Christella* clade species, populations and locations both within and outside Australia are required to be able to identify its parents.

The SNP assessment and hybrid phasing approach identified four additional putative hybrids in the dataset (Table 1). The three *Christella dentata* complex samples phased in a roughly 2:1 ratio to the *Amblovenatum queenslandicum* and *Christella subpubescens* references respectively, both in the *Christella* clade. This ratio might indicate second generation (F2) or later hybrids, or the result of a backcross with one parent. Both *Abacopteris aspera* also had high heterozygosity and allele divergence, but their reads phased to only one clade reference. This could indicate ancient hybrid origin of *Abacopteris aspera*, or introgression of alleles as a result of the hybridisation event that gave rise to the Russell River Fern and subsequent backcrossing. Three samples had high heterozygosity and allele divergence but did not phase in high proportion to any clade other than themselves: *Grypothrix triphylla* Almeida 4471, *Reholttumia truncata* Chong 158 and *Reholttumia sogerensis* Almeida 4477. Of these, *Grypothrix triphylla* is remarkable in having very high heterozygosity and allele divergence but only weak association to other clades (the highest being the *Chingia* clade, 9.9%, and *Plesioneuron* clade, 4.3%). This could indicate that *Grypothrix triphylla* is a hybrid between parents not sampled here, an ancient hybrid or the result of ghost introgression (Ottenburghs, 2020), or that its allele divergence results from paralogy due to polyploidy.

Replacing accessions with phased haplotypes accessions improved the phylogenetic reconstructions (Fig 5.). For example, the support values for the *Abacopteris* clade and the two sections of *Christella* were much higher than in the unphased phylogeny (Fig. 3). Split networks also showed that these sections were more clearly defined with fewer reticulations after phasing. This is strong evidence that phasing successfully reduced conflicting data in the analysis by phasing hybrid samples.

Our study successfully employed phasing of fern hybrids providing further proof-of-concept of the novel HybPhaser pipeline (Nauheimer et al., 2020). The combination of target sequence capture and hybrid phasing analysis give insights into the evolution of a lineage that were not possible in previous studies. The high proportion of species for which there is evidence of hybridisation in our Australian sample (24 % of samples) indicate hybridisation and reticulate evolution could be common and should be factored into studies in Thelypteridaceae globally. We expect hybridisation to be of similar importance in many other groups of ferns and believe utilisation of a hybrid phasing method such as HybPhaser should become routine in phylogenomic inference of fern target sequence capture datasets. We also expect that as evidence for the prevalence of hybridisation mounts for plants in general, the unilateral representation of evolution as a bifurcating tree, and especially the in-use taxonomic nomenclature framework will need to be revised.

## Acknowledgments

Data Accessibility Statement: Archival location will be provided upon acceptance. Collections were made by authorised officer Ashley Field, Queensland Herbarium, Department of Environment and Sciences. We thank Robert Jago and Nada Sankowsky for their assistance with sampling, Mel Harrison and Lalita Simpson for assistance with laboratory work and Jirrbal, Kuuku Ya’u, Ngadjon-ji and Yalanji Traditional Owners for access to traditional lands for sampling.

